# Detection of Pro-apoptotic BaxΔ2 Proteins in the Human Cerebellum

**DOI:** 10.1101/245910

**Authors:** Adriana Mañas, Aislinn Davis, Sydney Lamerand, Jialing Xiang

**Affiliations:** Department of Biology, Illinois Institute of Technology, Chicago, IL 60616

**Keywords:** BaxΔ2, Cerebellum, Purkinje Cells, Apoptosis, Human Brain

## Abstract

BaxΔ2 is a pro-apoptotic protein originally discovered in colon cancer patients with high microsatellite instability. Unlike most pro-apoptotic Bax family members, BaxΔ2 mediates cell death through a non-mitochondrial caspase 8-dependent pathway. In the scope of analyzing the distribution of BaxΔ2 expression in human tissues, we examined a panel of human brain samples. Here, we report 4 cerebellar cases in which the subjects had no neurological disorder or disease documented. We found BaxΔ2 positive cells scattered in all areas of the cerebellum, but most strikingly concentrated in Purkinje cell bodies and dendrites. Two out the four subjects tested had strong BaxΔ2- positive staining in nearly all Purkinje cells; one was mainly negative; and one had various levels of positive staining within the same sample. Further genetic analysis of the Purkinje cell layer, collected by microdissection from two subjects, showed that the samples contained G7 and G9 *Bax* microsatellite mutations. Both subjects were young and had no diseases reported at the time of death. As the distribution of BaxΔ2 is consistent with that known for Baxα, but in a less ubiquitous manner, these results may imply a potential function of BaxΔ2 in Purkinje cells.

## Introduction

The cerebellum is the area of the brain responsible for the modification of motor commands and is involved in balance, posture, voluntary movements, motor learning and more (Koziol et al. 2014; Bernard and Seidler 2014). The cerebellar cortex is composed of three layers, the granular layer, the Purkinje cell layer, and the molecular layer. The dendritic trees from Purkinje cells extend into the molecular layer (Casoni et al. 2017). Recent studies showed that Purkinje cells play a role in information storage for learned response sequences in coordination of motor behaviors (Jirenhed et al. 2017). It also has been shown that Purkinje cells may be involved in the pathophysiology of schizophrenia (Picard et al. 2007; Mothersill et al. 2016).

Bax is a pro-apoptotic Bcl-2 family member, that is ubiquitously distributed throughout all human tissues (Krajewski et al. 1994; Penault-Llorca et al. 1998). Bax plays a critical role in development and in tissue homeostasis, and its dysregulation can lead to many diseases. Due its involvement in the development and progression of neurodegenerative diseases and ischemic insults, expression and distribution of Bax in the brain has been widely studied (Hara et al. 1996; Fan et al. 2001; Vogel 2002; Dorszewska et al. 2004; Jung et al. 2008; Didonna et al. 2012; Garcia et al. 2013). Bax is known to have several functional isoforms, but only the distribution of the canonical isoform, Baxα, has been fully studied. Baxα is present in all the different areas of the brain, mainly in neuronal bodies, with very low to no presence in glial cells (Hara et al. 1996; Vogel 2002; Didonna et al. 2012). Cerebellar Purkinje cells and hippocampal neurons have the highest levels of Baxα (Hara et al. 1996; Vogel 2002; Casoni et al. 2017), which is believed to be the reason why these cells are so vulnerable to ischemic insults (Krajewski et al. 1995). Apart from its apoptotic role, other physiological functions of Baxα in the brain are unknown.

BaxΔ2 is a unique isoform of the Bax subfamily originally discovered in colorectal cancer patients with high microsatellite instability (MSI-H) (Haferkamp et al. 2012; Zhang et al. 2015). It is generally believed that generation of BaxΔ2 requires a microsatellite frameshift mutation in combination with an alternative splicing event that restores the reading frame. Like Baxα, BaxΔ2 is pro-apoptotic and has similar characteristics, such as binding with Bcl-2; however, BaxΔ2 does not target mitochondria, and instead activates a caspase 8-dependent death pathway (Haferkamp et al. 2012; Zhang et al. 2014; Mañas et al. 2017). In the process of analyzing the expression and distribution of BaxΔ2 throughout the human body, we examined several tissue sections of human brain. Here, we report the expression and distribution of BaxΔ2 in the cerebellum of four young and healthy human subjects.

## Materials and methods

### Materials

All tissue sections and tissue microarray slides were commercially obtained from Biomax. All samples were de-identified and assigned with codes. Antibody against BaxΔ2 was generated previously (Haferkamp et al. 2012).

### Immunohistochemistry and tissue analysis

Tissue slides were de-waxed using xylene and rehydrated using graded ethanol solutions. Endogenous peroxidase activity was blocked using 0.3% hydrogen peroxide. Slides were incubated in sodium citrate buffer (0.01 M, pH 6.0) at 95°C for epitope retrieval. After blocking with 5% BSA, CoverWell^®^ incubation humidity chambers were used to incubate the slides with anti-BaxΔ2 antibody (1:100 in blocking buffer) at 4°C overnight and then with Biotin-conjugated Goat anti-mouse secondary antibody (1:200 in PBS) at room temperature for 2 hours. Vectrastain^®^ ABC Kit (Vector Laboratories) and ImmPACT™ DAB Peroxidase Substrate Kit (Vector Laboratories) were used for visualization, and Hematoxolin QS (Vector Laboratories) was used for nuclear staining. Finally, slides were dehydrated and fixed using xylene based mounting media Poly-Mount^®^ (Polysciences Inc.). Fluorescence staining was performed as described for the DAB staining until the step of the primary antibody incubation. Slides were then incubated with Alexa Fluor^®^ 488 donkey anti-mouse IgG (Invitrogen) [1:200] at room temperature for 2 hours. Slides were fixed using ProLong^®^ Gold antifade reagent (Invitrogen). Slides were scanned at the Integrated Light Microscopy Core Facility at the University of Chicago and visualized using Pannoramic Viewer 1.15.2. Each slide was analyzed by three independent viewers and samples with debated evaluation were analyzed by a fourth individual. Each sample was assigned an H-score value based on the intensity and number of positive cells.

### Microdissection and Genotyping

The stained tissue sections of cerebellum from Subjects 1 and 3 were uncovered using xylene and dehydrated with graded ethanol solutions. The Purkinje cells layer was microdissected under a microscope using a sterile needle. The tissue was collected and lysed with Proteinase K. DNA was then isolated using AMPure XP magnetic beads (Beckman Coulter). Sanger sequencing was performed at the DNA Sequencing & Genotyping Facility at the University of Chicago Comprehensive Cancer Center.

## Results and Discussion

The monomeric form of Baxα is ubiquitously distributed throughout different human tissues, including the brain (Hara et al. 1996). However, distribution of other Bax isoforms is unknown. In the process of screening the tissue distribution of BaxΔ2, we found 4 interesting cases involving brain cerebellar tissues. The patient information about these 4 subjects was summarized in Figure 1A. There were two males and two females, all between the ages of 15 and 35. All subjects were healthy at the time of death and had no tumors or neuronal diseases reported. All tissues were immunohistochemically stained with an anti-BaxΔ2 antibody, which has been shown previously to be very specific for BaxΔ2 and has no cross-staining with parental Baxα. We detected significant amounts of BaxΔ2-positive cells in three out of the four subjects. As shown in Figure 1B, the most significant BaxΔ2-positive cells were found in the Purkinje cell layer, between the granular and molecular layers. Subjects 3 and 4 had strong BaxΔ2 staining in almost all Purkinje cells; Subject 2, on the other hand, had almost no positive staining observed; and Subject 1 had mixed populations of both positive and negative BaxΔ2 cells in close regions.

**Fig. 1.**
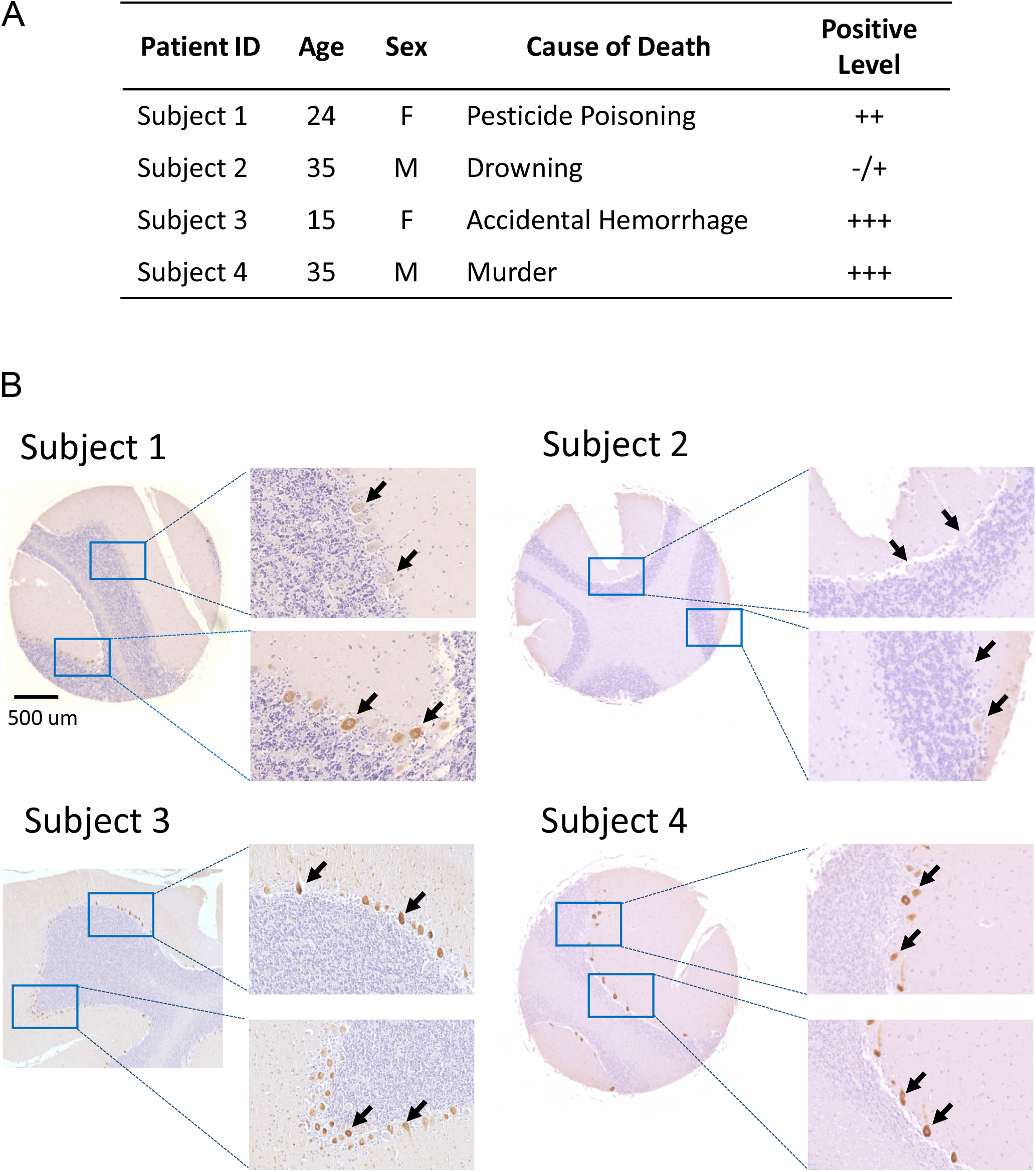
BaxΔ2 is expressed at different levels in the cerebella of four different subjects. A. Table summarizing the information of four subjects. B. Immunohistochemical staining of tissue sections with anti-BaxΔ2 antibody. The tissue sections show cerebella from the four subjects in (A). The non-enlarged tissue samples are presented at the same scale, scale bar 500 μm. Black arrows point at Purkinje cells presenting various levels of BaxΔ2 positive

Further analysis of the positive samples showed that most BaxΔ2-positive staining was detected in the Purkinje cell bodies (Fig. 2A), as well as in the dendrites extending into the molecular layer (Fig. 2D, 2E and 2F). In contrast with the strong positive-stained Purkinje cells, we also observed some weaker positive cells scattered throughout the granular layer (Fig. 2B) and the molecular layer (Fig. 2C). The nature of these cells remains to be determined; however, morphologically, they appeared to be granule cells and Golgi cells in the granular layer, and basket cells (large nuclei and close to the Purkinje cell layer) and stellate cells (smaller nuclei and body size) in the molecular layer. Overall, BaxΔ2 seemed evenly distributed in the cerebellum at low levels, with significantly higher BaxΔ2-positive staining in the Purkinje cells. These results are consistent with published data for the distribution of Baxα, which also presents higher levels in Purkinje cells. However, the amount and distribution of BaxΔ2- positive cells, outside of the Purkinje cell layer, was lower and less ubiquitous than for Baxα (Hara et al. 1996).

**Fig. 2.**
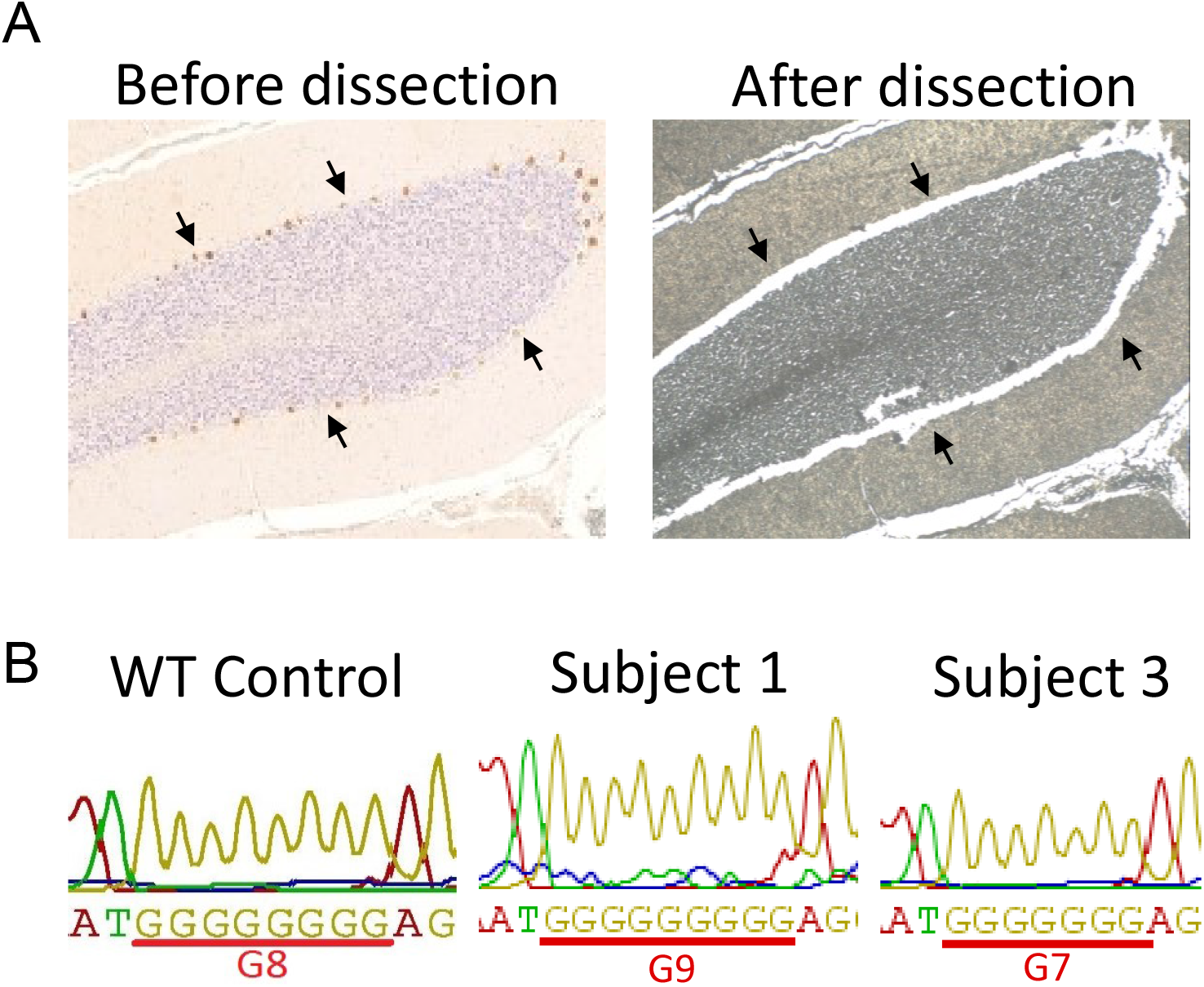
Positive BaxΔ2 cells are observed in all layers of the cerebellum. A. Immunohistochemical staining of the Purkinje cell layer from subject 1 with an enlarged BaxΔ2-positive Purkinje cell. B. Granular layer from Subject 3 with BaxΔ2 positive cells. C. Molecular layer from Subject 3 with several BaxΔ2-positive cells. D. Purkinje cell layer and molecular layer from Subject 1 with BaxΔ2-positive Purkinje cells. Black arrows point to BaxΔ2-positive dendrites. E and F. Fluorescence immunostaining of cerebellum sections from Subject 3 presenting BaxΔ2-positive staining in Purkinje cell bodies and dendrites. Red arrows point to BaxΔ2-positive dendrites

BaxΔ2 was previously detected in MSI-H tumors which involve mutations in the *Bax* microsatellite (Haferkamp et al. 2012). As none of these subjects were reported to have any tumors or neurological conditions, we wondered whether they could have a *Bax* microsatellite mutation. Due to limited sample size and variation of sample quality, we were only able to genetically analyze two of the four subjects. The tissue slides were first immunohistochemically stained with anti-BaxΔ2 antibody, then the Purkinje cell layer was carefully harvested through microdissection under a microscope, as shown in Figure 3A, and finally genomic DNA was isolated for sequence analysis. A 200 base-pair segment of the *Bax* gene containing the microsatellite region, was amplified by PCR and subjected to Sanger sequence analysis. As shown in Fig 3B, in comparison with the wild type *Bax* microsatellite, which contains a stretch of eight guanines (G8), the sample from Subject 1 contains a G9 mutation and the sample from Subject 3 contains a G7 mutation.

**Fig. 3.**
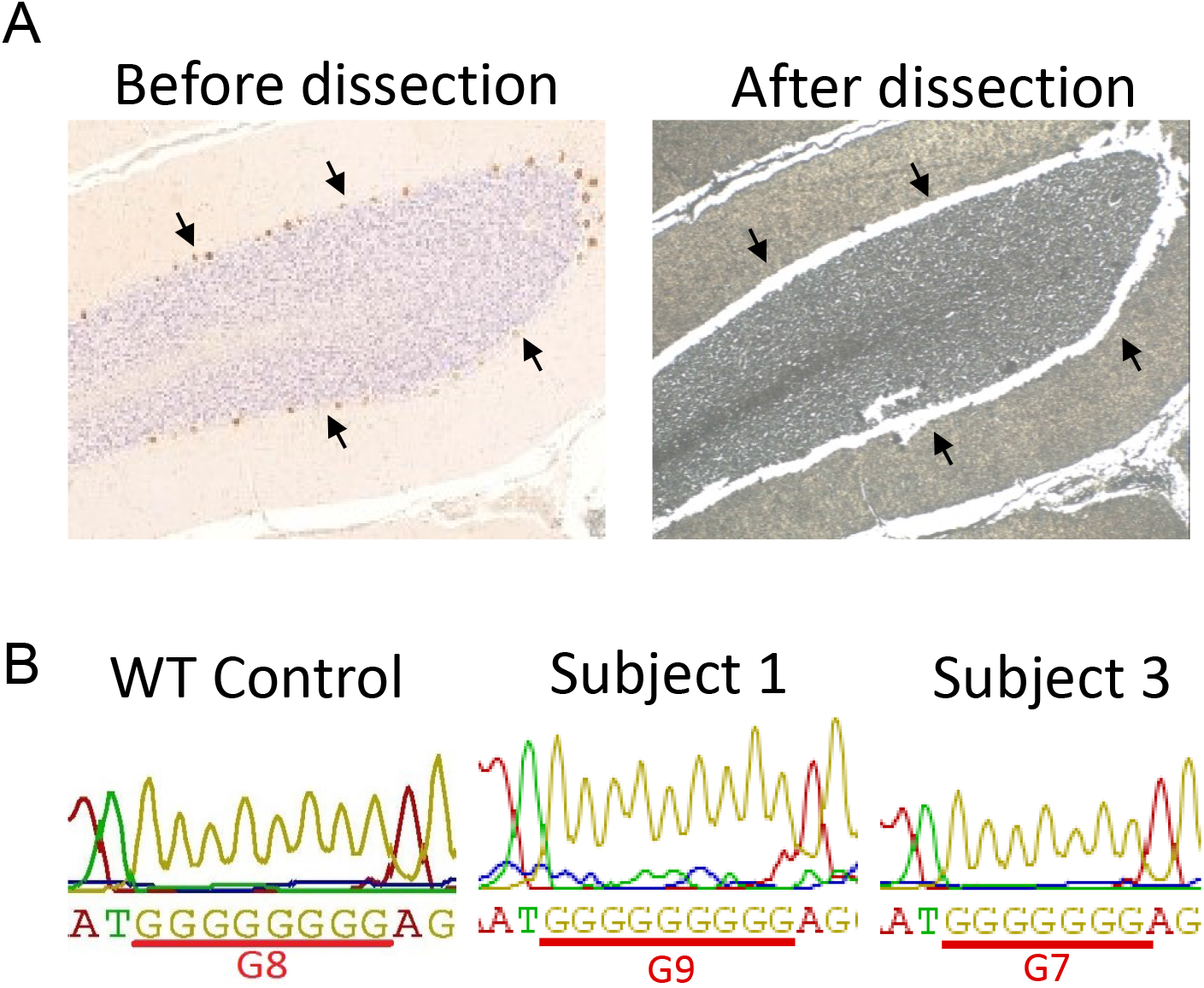
Genetic analysis of the BaxΔ2-positive Purkinje cell layer from two subjects shows mutations in the *Bax* microsatellite. A. Immunohistochemical staining of a cerebellum section from Subject 1, before and after microdissection of the Purkinje cell layer. Black arrows point to the Purkinje cell layer area dissected. B. Captions of Sanger sequencing data from the *Bax* microsatellite region of a wild type (G8) control, subject 1 (G9), and subject 3 (G7)

It is generally believed that expression of BaxΔ2 requires a *Bax* microsatellite G7 mutation and an alternative splicing event, which salvages the frameshift caused by the guanine deletion (Haferkamp et al. 2012). The result from Subject 3 seemed consistent with the BaxΔ2-positive staining, as the G7 mutation was detected. However, a G9 mutation was detected for Subject 1, who also had a strong BaxΔ2 staining in the area dissected, similar to that in Subject 3. This cannot be explained by the general rule for BaxΔ2 expression. Although we have previously shown that the G9 mutation can generate another BaxΔ2 sub-isoform, BaxΔ2(G9), which has a similar behavior as BaxΔ2 (Haferkamp et al. 2013), the antibody used here is specific for BaxΔ2 and does not cross react with BaxΔ2(G9). One possibility is that the area collected by microdissection also contained a significant amount of unstained cells with predominant G9 genotype. Another possibility is that the G8-to-G7 deletion could occur at the transcriptional level by transcriptional slippage, or at the translational level by ribosomal frameshift (Ketteler 2012; Atkins et al. 2016). Both transcriptional slippage and ribosomal frameshift are currently under investigation in our laboratory.

Nevertheless, we report the discovery of pro-apoptotic BaxΔ2 proteins in cerebellar neuron cells of young healthy individuals. The similar distribution pattern as canonical Baxa, especially in Purkinje cells, implies a potential physiological function of BaxΔ2 in neurons, independently or in compensation of Baxα.

## Acknowledgements and Funding

This work was supported by the National Institutes of Health [R15 CA195526].

## Compliance with Ethical Standards

The authors declare they have no conflict of interest.

## References

Atkins JF, Loughran G, Bhatt PR, Firth AE, Baranov P V. (2016) Ribosomal frameshifting and transcriptional slippage: From genetic steganography and cryptography to adventitious use. Nucleic Acids Res 44:gkw530. doi:10.1093/nar/gkw530

Bernard JA, Seidler RD (2014) Moving forward: Age effects on the cerebellum underlie cognitive and motor declines. Neurosci Biobehav Rev 42:193-207. doi:10.1016/J.NEUBIOREV.2014.02.011

Casoni F, Croci L, Cremona O, Hawkes R, Consalez GG (2017) Early Purkinje Cell Development and the Origins of Cerebellar Patterning. In: Development of the Cerebellum from Molecular Aspects to Diseases. Springer International Publishing, Cham, pp 67-86

Didonna A, Sussman J, Benetti F, Legname G (2012) The role of Bax and caspase-3 in doppel-induced apoptosis of cerebellar granule cells. Prion 6:309-316. doi:10.4161/pri.20026

Dorszewska J, Adamczewska-Goncerzewicz Z, Szczech J (2004) Apoptotic proteins in the course of aging of central nervous system in the rat. Respir Physiol Neurobiol 139:145-155. doi:10.1016/J.RESP.2003.10.009

Fan H, Favero M, Vogel MW (2001) Elimination of Bax Expression in Mice Increases Cerebellar Purkinje Cell Numbers but Not the Number of Granule Cells. 91:82-91

Garcia I, Crowther AJ, Gama V, Ryan Miller C, Deshmukh M, Gershon TR (2013) Bax deficiency prolongs cerebellar neurogenesis, accelerates medulloblastoma formation and paradoxically increases both malignancy and differentiation. Oncogene 32:2304-2314. doi:10.1038/onc.2012.248

Haferkamp B, Zhang H, Kissinger S, Wang X, Lin Y, Schultz M, Xiang J (2013) BaxΔ2 Family Alternative Splicing Salvages Bax Microsatellite-Frameshift Mutations. Genes and Cancer 4:501-512. doi:10.1177/1947601913515906

Haferkamp B, Zhang H, Lin Y, Yeap X, Bunce A, Sharpe J, Xiang J (2012) BaxΔ2 Is a Novel Bax Isoform Unique to Microsatellite Unstable Tumors. J Biol Chem 287:34722-34729. doi:10.1074/jbc.M112.374785

Hara A, Hirose Y, Wang A, Yoshimi N, Tanaka T, Mori H (1996) Localization of Bax and Bcl-2 proteins, regulators of programmed cell death, in the human central nervous system. Virchows Arch 429-429:249-253. doi:10.1007/BF00198341

Jirenhed D-A, Rasmussen A, Johansson F, Hesslow G (2017) Learned response sequences in cerebellar Purkinje cells. Proc Natl Acad Sci U S A 114:6127-6132. doi:10.1073/pnas.1621132114

Jung A-R, Kim TW, Rhyu IJ, Kim H, Lee YD, Vinsant S, Oppenheim RW, Sun W (2008) Misplacement of Purkinje cells during postnatal development in Bax knock-out mice: a novel role for programmed cell death in the nervous system? J Neurosci 28:2941-8. doi:10.1523/JNEUROSCI.3897-07.2008

Ketteler R (2012) On programmed ribosomal frameshifting: the alternative proteomes. Front Genet 3:242. doi:10.3389/fgene.2012.00242

Koziol LF, Budding D, Andreasen N, D’Arrigo S, Bulgheroni S, Imamizu H, Ito M, Manto M, Marvel C, Parker K, Pezzulo G, Ramnani N, Riva D, Schmahmann J, Vandervert L, Yamazaki T (2014) Consensus Paper: The Cerebellum’s Role in Movement and Cognition. The Cerebellum 13:151-177. doi:10.1007/s12311-013-0511-x

Krajewski S, Krajewska M, Shabaik A, Miyashita T, Wang HG, Reed JC (1994) Immunohistochemical determination of in vivo distribution of Bax, a dominant inhibitor of Bcl-2. Am J Pathol 145:1323-36

Krajewski S, Mai JK, Krajewska M, Sikorska M, Mossakowski MJ, Reed JC (1995) Upregulation of bax protein levels in neurons following cerebral ischemia. J Neurosci 15:6364-76

Mañas A, Wang S, Nelson A, Li J, Zhao Y, Zhang H, Davis A, Xie B, Maltsev N, Xiang J (2017) The functional domains for BaxΔ2 aggregate-mediated caspase 8- dependent cell death. Exp Cell Res 359:342-355. doi:10.1016/J.YEXCR.2017.08.016

Mothersill O, Knee-Zaska C, Donohoe G (2016) Emotion and Theory of Mind in Schizophrenia—Investigating the Role of the Cerebellum. The Cerebellum 15:357-368. doi:10.1007/s12311-015-0696-2

Penault-Llorca F, Bouabdallan R, Devilard E, Charton-Bain M-C, Hassoun J, Birg F, Xerri L (1998) Analysis of BAX expression in human tissues using the Anti-BAX, 4F11 monoclonal antibody on paraffin sections. Pathol - Res Pract 194:457-464. doi:10.1016/S0344-0338(98)80114-3

Picard H, Amado I, Mouchet-Mages S, Olie J-P, Krebs M-O (2007) The Role of the Cerebellum in Schizophrenia: an Update of Clinical, Cognitive, and Functional Evidences. Schizophr Bull 34:155-172. doi:10.1093/schbul/sbm049

Vogel MW (2002) Cell death, Bcl-2, Bax, and the cerebellum. The Cerebellum 1:277-287. doi:10.1080/147342202320883588

Zhang H, Lin Y, Manas A, Zhao Y, Denning MF, Ma L, Xiang J (2014) BaxDelta2 Promotes Apoptosis through Caspase-8 Activation in Microsatellite-Unstable Colon Cancer. Mol Cancer Res 12:1225-1232. doi:10.1158/1541-7786.MCR-14-0162

Zhang H, Tassone C, Lin N, Mañas A, Zhao Y, Xiang J (2015) Detection of Bax Microsatellite Mutations and BaxDelta2 Isoform in Human Buccal Cells. J Cell Sci Ther s8:. doi:10.4172/2157-7013.S8-002

